# Otters, but not apes, prepare for mutually exclusive possibilities

**DOI:** 10.64898/2026.02.03.703448

**Authors:** Sara Torres Ortiz, Kasper Fjorside, Paula Fuentes Raigal, Rocio Canales, Magnus Wahlberg

## Abstract

The ability to prepare for mutually exclusive outcomes is often considered uniquely human. Solving such problems requires anticipating alternative futures before acting. In the classic forked-tube task, the optimal strategy is to block both exits to secure a reward: children under four years and great apes typically fail, whereas older children succeed. Using this paradigm, we tested three Asian small-clawed otters (*Aonyx cinereus*) and one Eurasian otter (*Lutra lutra*) and compared their performance with chimpanzee data. Otters covered both exits significantly more often than chimpanzees, with all individuals succeeding within their first trials. Initial inconsistency in maintaining the strategy appeared linked to anatomical constraints that limited reward success. When retested two months later with an apparatus better suited to otter morphology, individuals adopted and maintained dual coverage as success increased, indicating that the behavior tracked the payoff structure of the task rather than reflecting low-level mechanisms such as trial-and-error learning. Together, these findings indicate that blocking both exits is an adaptive response to the task’s causal structure, supporting the ecological intelligence hypothesis: cognition evolves in response to ecological demands, particularly foraging challenges that place recurrent pressures on memory, decision-making, and executive control, rather than being driven solely by social complexity.

## Introduction

The ability to prepare for uncertain futures is a distinguishing characteristic of human cognition, requiring not only anticipation of a single event but also the ability to represent multiple, mutually exclusive possible outcomes [1,2]. For example, when crossing a street, humans instinctively look both ways, preparing for two alternative futures. Developmental research indicates that this ability emerges relatively late in children, typically between 3 and 4 years of age, and is not reliably demonstrated in non-human apes [3,4] even though such problems are common in nature: an opening that allows prey to escape cannot simultaneously remain blocked. Solving these tasks efficiently requires more than associative learning; it requires anticipating exclusivity before acting, a capacity sometimes described as anticipatory reasoning [5]. The apparent limitation of apes in showing that capacity has been interpreted as evidence that abstract reasoning about exclusivity evolved late and may depend on executive capacities associated with large brains or language precursors.

A growing body of research challenges this anthropocentric view. Increasingly, sophisticated cognitive strategies have been identified in distantly related species with very different neural architectures. Cleaner wrasses adjust their behavior strategically when reputational consequences are at stake [6]; cuttlefish optimize foraging sequences using episodic-like memory [7]; and bees and fish can perform inference-by-exclusion and quantity discrimination tasks once thought to only be solvable by primates [8–10]. These discoveries point toward a more general principle name the ecological intelligence hypothesis: complex cognition emerges wherever ecological conditions reward structure-sensitive decision making, not only within large-brained or socially complex lineages [11].

Following this logic, the ability to anticipate mutually exclusive outcomes should evolve in species that frequently confront object-bounded problems with irreversible loss, for example, when prey can escape through multiple openings or when manipulation of one path constrains another. Such conditions characterize the foraging ecology of otters differing from apes where their usual diet relies in stationary items such as fruit and grains. Otters are highly dexterous carnivores that hunt fish, crustaceans, and mollusks in spatially complex environments such as rocky crevices or shells with multiple apertures. They routinely use their forepaws to probe, contain, and extract prey that might otherwise flee [12,13]. Several species also exhibit habitual tool use, manipulating stones or shells as hammers and anvils to open hard-shelled food [12]. These behaviors suggest that otters operate in an ecological niche where anticipatory behavior confers direct fitness benefits and where the motor demands of simultaneously controlling multiple apertures are well within their everyday manipulative repertoire.

The present study tests whether two species of otters, one social and one nonsocial, exhibit anticipatory reasoning when faced with a novel physical problem that embodies mutual exclusivity. We used the forked-tube paradigm in which a food item is dropped into a tube with two possible exits. The optimal strategy is to cover both openings, ensuring success regardless of outcome. Solving the task efficiently required recognizing that the two routes were mutually exclusive: securing one exit should preclude loss through the other. We predicted that if otters understand this exclusivity relation, they would block both exits before attempting to access the reward, thereby constraining the problem space prior to manipulation. By contrast, if they relied on trial-and-error exploration, they would interact with the tube or one exit first and only cover the second after observing food movement or failure as the apes did.

## Materials and Methods

### Experimental subjects

We tested three Asian small-clawed otters (*Aonyx cinereus*): two females (F1 and F2, both born 26 July 2021) and one male (M1, born 3 October 2009), as well as one female Eurasian otter (*Lutra lutra*; F3). The two Asian small-clawed females were captive-born at Zoo Dortmund (Germany) and transferred to Mundomar (Spain) on 9 June 2022, while the male was born at Mundomar.

The Eurasian otter was wild-born and rescued as an orphan from a construction site in Jutland, Denmark, in spring 2022. Estimated to be approximately one month old at intake, she was hand-reared at AQUA Aquarium & Wildlife Park (Silkeborg) before being transferred to the Marine Biological Research Center, University of Southern Denmark (Kerteminde), in early 2023.

All Asian small-clawed otters were housed in enclosures with indoor and outdoor areas. During the study, their primary enclosure was under renovation, and all training and testing occurred in a temporary quarantine facility comprising two connected areas (8 m^2^ and 25 m^2^) with a small pool (4 m long, 45 cm deep) allowing voluntary water access. Animals were expected to return to their permanent enclosure after data collection.

The Eurasian otter was housed in a 65 m^2^ enclosure with a 20 m^2^ (26 m^3^) pool and had daily access to both land and water, along with multiple daily training or enrichment sessions.

None of the otters had prior experience with cognitive testing or tube-based apparatus. The Eurasian otter had previous operant conditioning experience for psychophysical tasks.

### Diet and Experimental Reinforcement

Asian small-clawed otters were fed primarily freshwater fish (trout and tilapia fillets), which also served as experimental rewards. Additional items included whole carp, smelt (preferred by the male), mollusks (typically with shells), renal-support dry food (Hill’s Prescription Diet k/d for cats), and insects for enrichment. Reinforcement during trials consisted exclusively of small fish pieces.

The Eurasian otter received common roach (*Rutilus rutilus*) during training and was additionally fed chicken and small amounts of sprat daily. During testing, rewards consisted of small pieces of sprat or trout. Eggs, insects, and other items were occasionally provided as enrichment, and vitamin supplementation was administered each morning. She typically participated in three to four daily sessions, with at least one dedicated to enrichment.

### Apparatus

All subjects were tested using a tube-based apparatus designed to examine anticipatory responses to mutually exclusive future outcomes. The original goal was to replicate the classical forked-tube paradigm [4], consisting of a single entrance and two exits. However, due to the small size of otter forepaws and the PVC diameters required for effective exit coverage, food frequently became lodged inside the tube. Following prior work in chimpanzees showing similar motor limitations [14], we adopted a modified design using two separate tubes while preserving the critical feature of outcome uncertainty.

Subjects were first trained that food would exit a PVC tube and would be lost if the exit was not covered. For Asian small-clawed otters, two tubes (32 mm length, 50 mm diameter) were positioned so subjects could access the lower openings. An opaque plexiglass panel (17 × 17 cm) occluded the entrance, preventing visual tracking of the food trajectory. The Eurasian otter was initially tested with the same apparatus. After the first session, her configuration was modified by adding a 45° elbow joint to each tube to facilitate interception while maintaining exit uncertainty.

After completing the two-tube condition, we constructed a professionally built forked-tube apparatus for the Asian small-clawed otters to reduce motor constraints and improve reward rates. The vertical entrance tube (50 mm diameter) connected to two lateral exit arms (32 mm diameter) diverging at 45°. An internal plexiglass divider allowed the experimenter to control the exit side. Total length from entrance to exits was 24 cm.

All sessions were recorded using a high-speed digital camera (240 fps, 1080p) positioned to capture forelimb movements and tube exits, enabling frame-by-frame analysis of response timing, limb placement, and interception success. Subjects were free to cover one exit, the other, both simultaneously, or neither; no physical constraints were imposed.

### Procedure

During both training and testing, the experimenter released the food item only after subjects placed their forepaws in a stable position. If food slipped despite correct positioning, no compensatory reward was provided, ensuring reinforcement remained contingent on successful interception.

#### Training Phase

All otters completed an initial training phase using a single PVC tube to establish that uncovered exits resulted in reward loss. The tube was positioned behind a fence allowing forepaw extension; uncovered food fell outside the enclosure.

Asian small-clawed otters began training on 14 August 2025. Individuals were trained simultaneously with one experimenter per animal. Sessions consisted of 12 trials with randomized, counterbalanced baiting locations, ensuring no side was baited more than twice consecutively. Each otter completed 96 trials before entering the test phase on 4 September 2025.

The Eurasian otter began training on 20 August 2025 under the same protocol, completing 96 trials before testing on 9 September 2025.

Prior to testing, subjects observed 10 demonstration trials of the two-tube configuration with exits out of reach, confirming that food emerged from only one exit per trial.

#### Test Phase: Two-Tube Condition

All two-tube testing occurred within a single day per subject. Asian small-clawed otters completed 72 trials across six sessions (12 trials each) with 30–90 min breaks. The Eurasian otter completed 96 trials across two sessions of 48 trials.

Trial order was randomized and baiting locations counterbalanced, with no side baited more than twice consecutively. At each trial, food was dropped into the occluded entrance. Subjects could cover the left exit, right exit, both exits, or neither. Successful coverage resulted in reward acquisition; otherwise, the food was lost. No corrective feedback was provided.

#### Test Phase: Forked-Tube Condition

Asian small-clawed otters were tested with the forked-tube apparatus on 27–28 October 2025, approximately two months after the two-tube tests. Due to frequent blockages, only one forked tube functioned reliably.

F1 completed 74 trials across four sessions; F2 completed 63 trials across four sessions over two days. M1 completed 42 trials across three sessions, but a mechanical failure during the first session produced a strong right-side bias. Although corrected later, his forked-tube data were excluded from analyses.

Trial order remained randomized and counterbalanced, with no visual access to the food trajectory.

### Behavioral Coding and Analysis

All trials were video-recorded and analyzed offline using slow-motion playback (240 fps). Responses were coded relative to the moment of food release, recording whether subjects covered the left exit, right exit, both exits, or neither. Covering behavior required physical occlusion preventing unobstructed passage of the food.

Reward outcome (obtained vs. lost) was recorded for each trial. For response stability analyses, run lengths were extracted as the maximum number of consecutive dual-coverage trials.

Analyses were conducted separately for the two-tube and forked-tube tasks. Otter performance in the two-tube task was compared with previously published chimpanzee data using independent-samples t-tests on proportions of dual and single-exit coverage, with adjusted degrees of freedom where necessary. Changes in individual strategy use across tasks were assessed using chi-square tests. M1’s forked-tube data were excluded due to apparatus malfunction. All tests were two-tailed (α = 0.05), with individual-level analyses capturing inter-individual variability.

Reward rates were calculated as the proportion of trials resulting in food acquisition, both overall and conditional on response type. Associations between response and reward were assessed descriptively using conditional probabilities. Session-level response rates were visualized with binomial 95% confidence intervals (Wilson method).

All statistical analyses and figure generation were conducted in R [15]. Data wrangling and session-level summaries used dplyr [16], plotting used ggplot2 [17], and confidence intervals were computed programmatically using iteration utilities from purr [18]. Percentage axis labeling used scales. Data import from Excel used readxl [19].

## Results

Three Asian small-clawed otters (*Aonyx cinereus*; F1, age 4; F2, age 4 and M1, age 16) and one Eurasian otter (*Lutra lutra*; F3, age 3) participated in this experiment. Because fish frequently became stuck inside forked tubes and otter forelimb morphology makes simultaneous control of two narrow branches, we first used a modified two-tube apparatus with concealed upper openings (Two-tube task; Figure 1C) preserving uncertainty while eliminating mechanical obstructions. Across the 96 trials conducted in a single session on Day 1, otters covered both exits significantly more often than chimpanzees tested in previous studies (4, 14; percentage of trials with dual coverage: t(12.4) = –2.8, P = 0.015; Figure 1B). In contrast, there were difference between otters and chimpanzees in the proportion of trials in which subjects covered only the right exit (t(6.0) = 1.5, P = 0.18) or only the left exit (t(6.7) = –0.69, P = 0.51). Despite their higher rate of dual coverage, no otter sustained this strategy consistently, and overall reward rates remained low (48.6%), likely reflecting anatomical constraints of the otter forelimb, including limited wrist rotation that hampers simultaneous control of two apertures.

**Figure 1.**
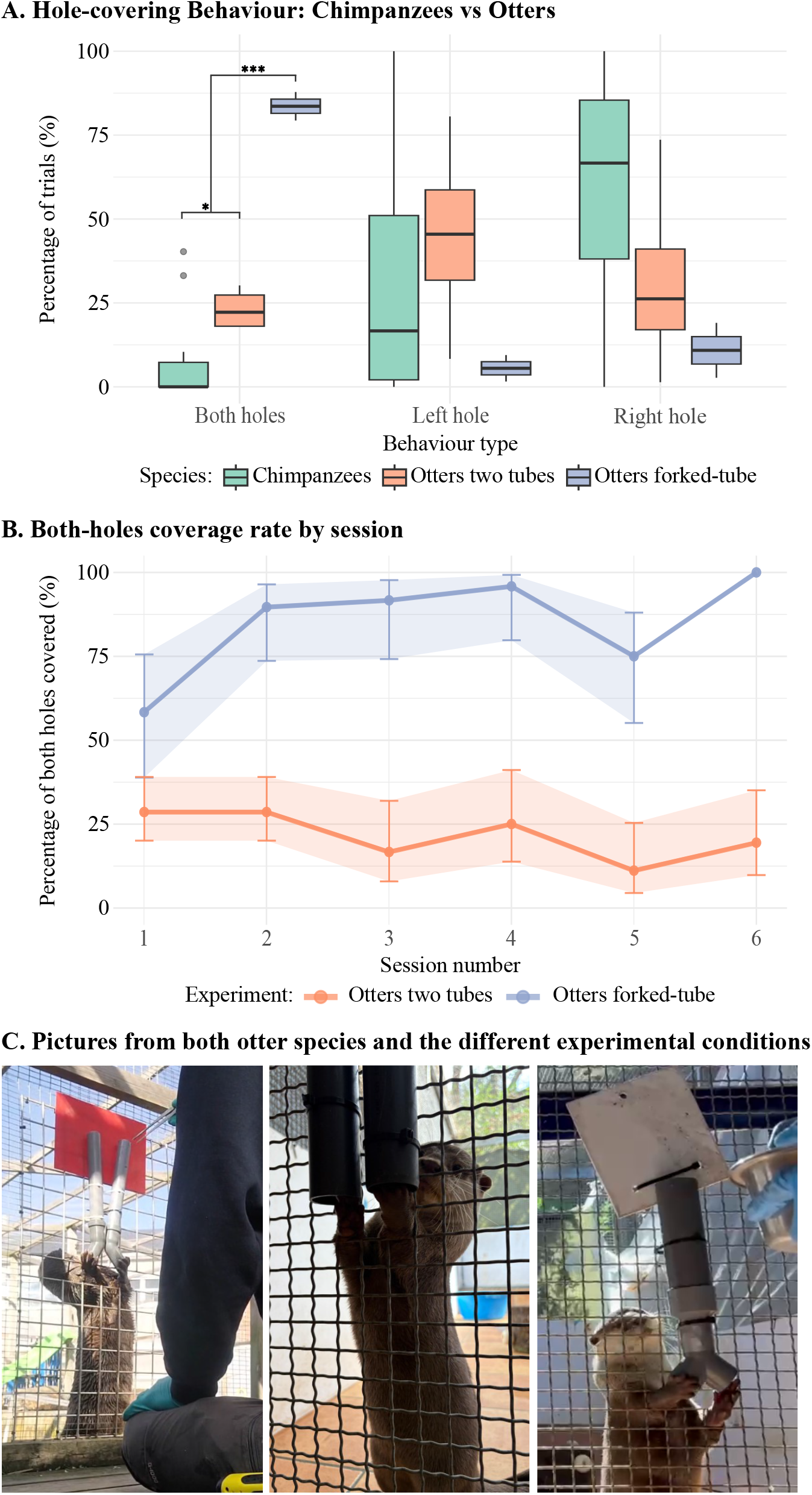
Experimental results and testing procedure. **(A)** Boxplot showing the number of trials in which the left hole, right hole, or both holes were covered, combining the otter data in both experimental conditions with chimpanzee datasets from Redshaw and Suddendorf (4) and Lambert and Osvath (14). **(B)** Proportion of trials with both openings covered across sessions in the Two-tubes and Forked-tube experiments. Each point shows the session-level proportion of trials in which an otter covered both available openings, plotted separately for the Two-tubes (orange) and Forked-tube (blue) conditions. Lines connect successive sessions within each experiment. Vertical error bars and shaded ribbons indicate 95% binomial confidence intervals for the session means, reflecting uncertainty due to the number of trials in each session. **(C)** Example trial from both otter species in both experimental conditions in which individuals successfully covered both holes.

Dual coverage emerged rapidly: F1 in her first four trials, F2 in trial 3, F3 in trial 4, and M1 in trial 2. To further reduce motor constraints, a redesigned forked-tube was introduced 50 days later, in a second single-day test session (Day 2; Figure 1C). All otters immediately covered both exits on their first trials with this device (Figure 1A).

Across full testing on Day 2 with the modified device, both females consistently adopted the “both” strategy (F1: 65/74 trials, 87.8%; F2: 50/63, 79.4%), whereas M1 did so rarely (5/42, 11.9%). M1’s performance was dominated by a persistent right-side bias triggered by an early tube malfunction. Even though the functionality of his tube was subsequently improved, his behavior did not change in the remaining trials: M1 chose the right exit in 36/42 trials (85.7%), producing a pronounced and consistent side bias that likely masked his ability to implement the anticipatory strategy. Relative to their performance in the two-tubes task, both females showed a significant increase in the adoption of the “both” strategy with the forked-tube (F1: χ^2^ = 72.4, P = 1.2×10□^1^□; F2: χ^2^ = 48.1, P = 3.6×10□^12^). Also, the reward rate when otters covered both exits was higher for the new apparatus (75% vs. 48.65% for the two-tubes). By contrast, M1 showed no reliable shift toward dual coverage (χ^2^ = 2.5, P = 0.11). Their behavior was temporally stable, with runs of 19 (F1) and 18 (F2) consecutive dual-coverage trials, whereas M1 showed no such stability (maximum run = 2).

## Discussion

Our findings show that otters spontaneously adopted the dual-coverage strategy within their first few trials and at substantially higher rates than chimpanzees tested in comparable studies [3,14]. This pattern emerged too early and too consistently to be explained by trial-and-error learning or indiscriminate blocking [4]. Instead, otters’ behavior tracked the informational structure of the task: dual coverage was used selectively when it was advantageous, sustained across long trial sequences, and increased specifically in the condition where the two future outcomes were mutually exclusive.

The reward structure of the task supports this interpretation. In the two-tubes condition, covering both openings conferred no payoff advantage because anatomical constraints prevented reliable interception. By contrast, in the forked-tube condition, dual coverage increased success to 75%, making it the only strategy that improved reward rate. This demonstrates that blocking both exits was not a habitual motor pattern but an adaptive response to the causal structure of the problem.

The divergence between our results and those observed in non-human primates [4,14] suggests that the cognitive requirements of representing bifurcating futures may be less constrained by impulsivity in specialized carnivores. In primate studies, failure is often attributed to a “pre-potent” urge to reach directly for the reward rather than executing a preparatory, indirect action [20]. Our otters, however, demonstrated a remarkable ability to inhibit direct pursuit in favor of a “split” motor plan. This high degree of inhibitory control may be rooted in the specialized neuroanatomy of the carnivore striatum. Recent volumetric assessments of large carnivores reveal a “hypertrophic” caudate nucleus, a structure critical for goal-directed decision-making and behavioral inhibition, that is disproportionately larger and more robustly connected to the frontal cortex than that of primates [21]. This suggests that the executive control required to prepare for mutually exclusive outcomes may have evolved through different selective pressures than those found in generalist primates.

Motor limitations also cannot explain the results. Restricted wrist rotation and difficulty maintaining bilateral contact reduced success rates but would tend to obscure, rather than produce, strategic dual coverage. When these constraints were removed in a modified apparatus, all otters immediately covered both exits. The male’s side bias, triggered by an early mechanical failure, further illustrates how apparatus-level factors can mask competence, a pattern well documented in primates and elephants on this task [3,14,22].

This performance contrasts sharply with non-human apes [3,14]. Chimpanzees typically fail to adopt dual coverage even after extensive experience, and elephants only rarely do so after hundreds of trials [22], consistent with incremental learning rather than anticipatory reasoning. In contrast, all otters succeeded immediately, and the two females maintained the strategy over extended runs in a condition where it yielded clear reward benefits, providing strong evidence that they anticipated mutually exclusive outcomes before acting.

Ecologically, this fits otter foraging biology and supports the ecological intelligence hypothesis [11,23]. Otters regularly pursue prey in crevices, shells, and other multi-aperture structures where loss through an unguarded exit is common, making advance constraints of alternatives highly advantageous [13,24–26].

That both a largely solitary species and a highly social one showed the same pattern indicates that this form of anticipatory reasoning does not depend on social complexity or large brain size but instead reflects a broader ecological principle: cognition evolves adaptively with ecological demands, particularly foraging problems that place recurrent pressures on memory, decision-making, and executive control, rather than being driven solely by social complexity [11].

Crucially, the Lutrinae subfamily offers a powerful, underutilized model to decouple the social and ecological drivers of varied cognitive abilities. By exhibiting a spectrum of social systems (solitary to gregarious), ecosystems (freshwater to marine), and manual dexterities, otters provide a “natural experiment” to test the origins of intelligence. Moving beyond primate-centric paradigms, the otter model allows us to determine whether complex cognition is a byproduct of social living or an adaptive response to the specific challenges of specialized foraging and multi-aperture prey pursuit. Together, these findings challenge the view that representing alternative futures is uniquely human or evolutionarily late. Otters prepared for mutually exclusive outcomes in a way not readily explained by learning or motor heuristics, suggesting that the capacity to anticipate multiple possible futures may be more widely distributed across mammals than previously assumed.

## Ethics

Experimental protocols were approved by the Animal Care Committee of Mundo Mar and complied with international and institutional guidelines. All procedures were conducted in accordance with the Guidelines for the ethical treatment of nonhuman animals in behavioral research and teaching (Association for the Study of Animal Behaviour, Animal Behaviour, Volume 195, January 2023, Pages I–XI). In accordance with the Spanish Animal Welfare Act 32/2007 of 7th November 2007, Preliminary Title, Article 3, the study was classified as non-animal experiment and did not require any approval from a relevant body. The experiments with the Eurasian otter were made under a permit from the Danish Animal Experiments Inspectorate (Ministry of Food, Agriculture and Fisheries of Denmark, permit nr. 2023-15-0201-01608). All training and testing procedures were based exclusively on positive reinforcement, and participation was voluntary at all times. No food, social, or environmental deprivation was used to motivate behavior, and their full daily diet was given regardless of participation.

## Data accessibility

Our data was live coded during testing and subsequently entered in an Excel file. The dataset contains individuals’ trial-by-trial responses in a tube task designed to evaluate preparation for uncertain outcomes. Variables are defined as follows: Experiment (experimental condition; e.g., two-tube vs. alternative setup), Session (set of trials conducted within a single testing block), Trial (trial number within each session), Animal (individual identity), Side (location of the baited tube: left or right), Response (animal’s behavioral response: left, right, or both), and Reward (binary outcome; 0 = no reward, 1 = reward obtained). The file therefore provides a complete record of individual decision-making across trials, allowing reconstruction of response strategies and success rates over time. Our data is available here:

https://github.com/Sharinade/Otters-but-not-apes-prepare-for-mutually-exclusive-possibilities

## Declaration of AI use

We have not used AI-assisted technologies in creating this article.

## Authors’ contributions

S.T.O.: conceptualization, data curation, formal analysis, investigation, methodology, project administration, resources, visualization, writing—original draft, writing—review and editing; K.F.: data curation, methodology, resources, writing—review and editing; P.F.R.: data curation, methodology, resources, writing—review and editing; R. C.: data curation, methodology, resources, writing—review and editing; M.W.: funding acquisition, methodology, project administration, resources, supervision, writing— review and editing.

All authors gave final approval for publication and agreed to be held accountable for the work performed therein.

## Acknowledgments

We thank the care takers of Mundo Mar for their dedication and motivation during the study, particularly to Cristina Pritchett, Carla Alvares and Manu Rodriguez. We also thank Thomas Suddendorf and Jonathan Redshaw for inspiring us for this study and being open to discussing the methodology.

